# Efficient online-group-screening designs for agent identification

**DOI:** 10.1101/220863

**Authors:** Tianxiang Gao, Omri Finkel, Jeff Dangl, Vladimir Jojic

**Affiliations:** Department of Computer Science, University of North Carolina, Chapel Hill, North Carolina, USA; Department of Biology, University of North Carolina, Chapel Hill, North Carolina, USA; Howard Hughes Medical Institute, Chevy Chase, Maryland, USA

**Keywords:** experiment design, important factor identification, pre-screening, feature selection, 1-bit compressed sensing, high-throughput screening

## Abstract

Identifying significant causal agents among a large number of candidates is challenging. When experimental resources are limited, exhaustively screening a large number of agents for the desired effect could incur a large cost and take a substantial amount of time. However, in many large scale experiments, such as high-throughput screening (HTS), the ratio of causal to non-causal agents is usually very low.

In this paper, we introduce a group-screening strategy to efficiently screen causal agents by grouping them into treatments. Our analysis shows that when a large number of candidates factors are screened and true agent percentage is very low (less than 1%), even in the worst case we could save up to 80% of the experiment runs. In the case where experiments span many rounds, we provide an online version of the group-screening that can determine the best strategy automatically based on the existing results. We applied this method to a real HTS experiment with 50,000 candidates that would require 9 rounds to finish in an exhaustive case. Our analysis showed that by applying the online-group-screening method, in the worst case, we can use 3 rounds and 19.7% (9828/50000) total tests to identify all the agents.

Finally, we show that with minor modifications, this framework extends to more complex agent discovery problems.

## 1 Introduction

Understanding the causes of phenotypic changes is one of the key goals of the biological research. Demonstrating that a particular candidate (e.g gene, mutant, bacterial strain) effects the change, provides evidence for a mechanistic relationship between a phenotype and a candidate. We will call such candidates “agents”. Candidates that have no effect on phenotypes are called “non-agents”. Screening designs examine the ability of each candidate to induce the change in phenotype [1]. We define “treatment” as a group of candidates that can be tested for phenotypic changes. The total time required to test a treatment from setting up to collecting results is called one “round”. Typically, multiple treatments can be tested in parallel in a single round. We use “efficiency” to represent the maximal possible number of treatments that can be tested per round. In this paper, we aim to provide a design method to minimize the number of total rounds needed for identifying all agents.

Generally, a naive method will exhaustively test all the candidates individually. However, when there are too many candidates to be tested when efficiency is low, a naive method will take many rounds to finish. In plant and animal experiments, efficiency is limited by factors such as greenhouse space or incubator/cage number. Furthermore, typical biology experiments can take days to weeks to finish. We summarize the duration for some experiments in Table 1. For high-throughput screening (HTS) experiments, efficiency is limited by machine/robot/human experimenter efficiency. For example, in the HTS experiment conducted by Ma *et al.* [2], only 40-60 96-well plates were analyzed in a single round. It took 9 rounds to test all 50,000 candidate compounds. According to a review from Macarron *et al.* [3], in many drug discovery companies, compound screening step can take up to 3 months to finish (for HTS of 1 million compounds). These limitations prevent biologists from exploring more candidates.

**Table 1.**
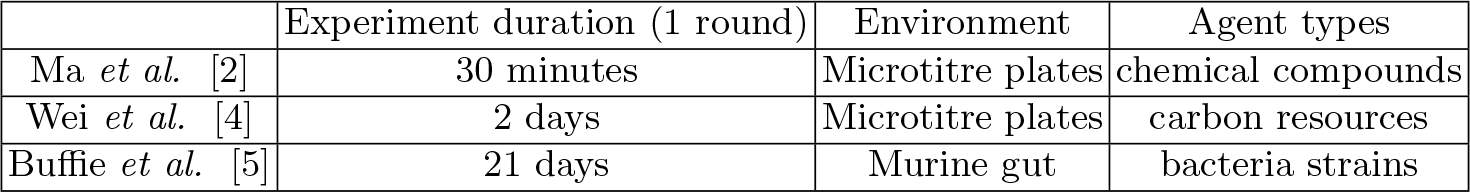
Example experiment durations

In many screening problems, true agents are actually very few, which is known as the “effect sparsity assumption” [6]. Using this assumption, it is possible to reduce the number of total treatments by applying methods like the one proposed by Gupta *et al.* [7]. They provided an adaptive sequential tree-searching algorithm for efficiently discovering relevant features in a binary classification problem requires *s* log *n* tests and *s* log *n* rounds, where *n* is the total number of features, and *s* is the number of relevant features. This algorithm can be recast in the language of agent identification. Features in this setting can be seen as candidates, samples as treatments required to be tested, and the binary label of a sample as the target phenotype. Thus collecting labeled samples is equivalent to measuring target phenotypes for the corresponding treatments. Loosely, the idea is to split the candidates into two treatments. If a treatment induces phenotypic change, the treatment is deduced to contain at least one agent, and the candidates in this treatment will be split further, recursively. If a treatment induces no phenotypic change, then none of its candidates are agents.

The drawbacks of this approach are: 1) it takes up to *s* log *n* rounds to identify all the agents. And in each round, it only tests one treatment, which does not fully utilize the experimental resources. For example, an experiment with 50,000 candidates and 7 agents will take more than 100 rounds to finish, no matter how many treatments we can test each round. Further, assays tested in a larger number of batches can exhibit “batch effects” [8]. 2) In most cases, we do not know the true agent number *s*. If the problem has *s* log *n > n*, this approach may result in much more treatments than the naive method. 3) In some cases, the number of candidates in one treatment can be limited because of the dilution limitation or maximal capacity of an assay.

In this paper, we present a group-screening method that can efficiently optimize the time required for discovery and overcome the drawbacks of the tree-searching method. This method subdivides *n* candidates into *b* group-screening treatments. According to the Pigeon-hole principle [9], whenever *b* is larger than the true agent number *s*, some treatments will not contain any true agents. Thus, we do not need to test the candidates in those treatments. We prove that we can identify all the agents using no more than 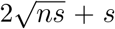 tests. We extend this method to **online-group-screening** that can utilize the results gathered at the end of each round to decide the best strategy based on a probabilistic analysis. Therefore, we can dynamically decide when to use group-screening and how many tests we can expect to save.

Figure 1 illustrates the group screening and online-group-screening methods. We have applied these methods to a real HTS experiment. Our results illustrate significant savings in both time and total tests. We believe this method will make agent identification problems less expensive, more efficient, and thus increase the pace of discovery.

**Fig. 1.**
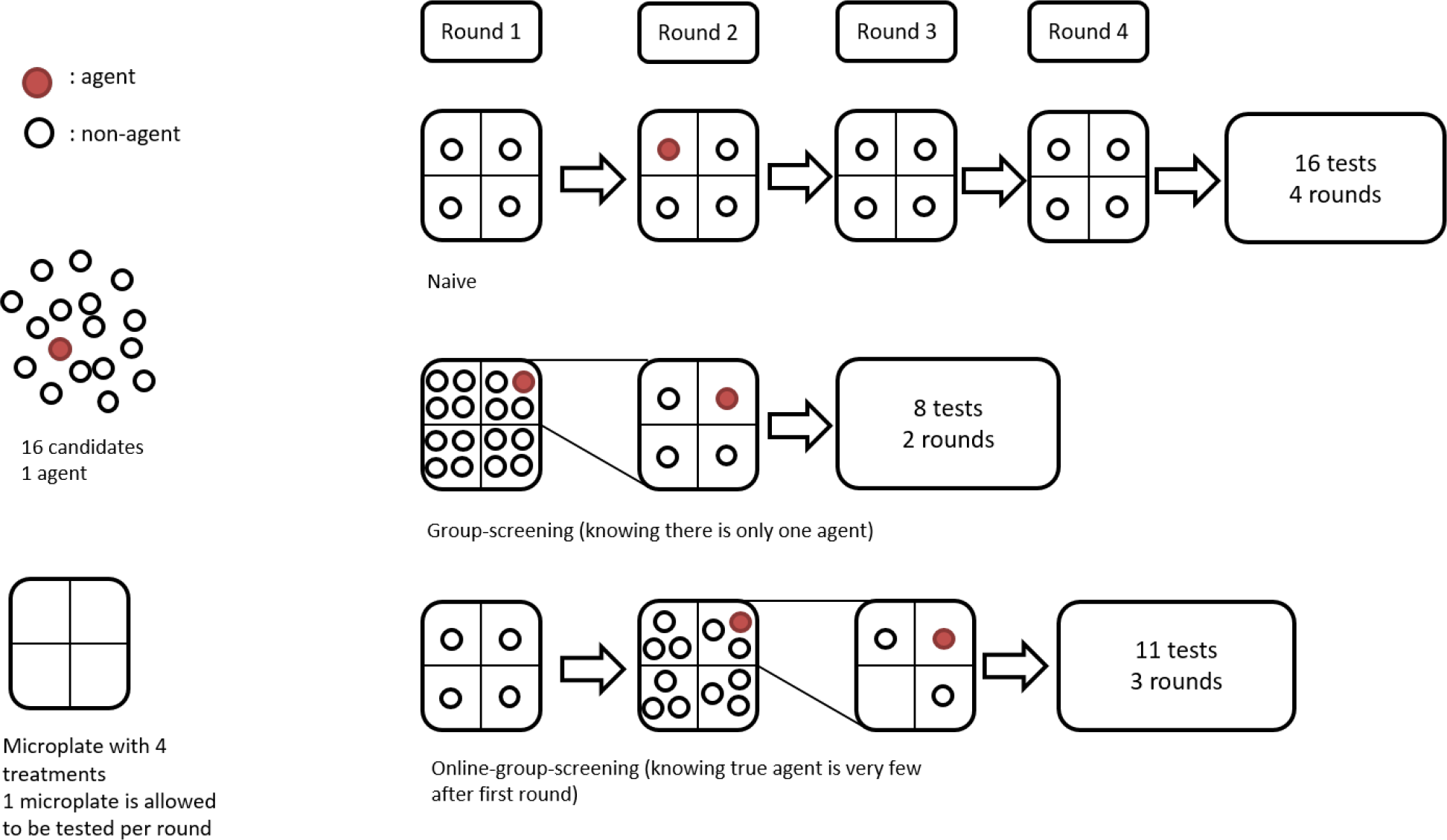
An example of naive, group-screening and online-group-screening algorithms. In this example, we have 16 candidates to test, one of which is a true agent. Each round we only allow at most 4 treatments to be tested in parallel, so efficiency is 4. The **naive** method exhaustively tests all candidates, using 16 tests and 4 rounds. If we know there is only one agent, the **group-screening** method randomly divides all 16 candidates into 4 group-screening treatments. After the first round, the treatment containing a true agent is further tested in the second round, using totally 8 tests in 2 rounds. When the true agent number is not known, we use the **online-group-screening** method. We run the first round as the naive method, and the results indicate that the true agent ratio is very low. Therefore, we use group-screening on the rest of the candidates. It takes 11 tests and 3 rounds in total.

In Section 2, we introduce our method and show its analytical and empirical performance. A case study of this method on a real HTS experiment is presented in Section 3. In Section 4, we illustrate how this framework can be extended to solve several different problems. Final discussion is in Section 5.

## 2 Method

In this section, we first specify the problem of agent identification. Next, we introduce the group-screening method and its theoretical savings compared to the naive method. Finally, we will present an online version that can adaptively choose the best method.

### 2.1 Problem specification

We assume that there are *n* candidates *U* = *{x*_1_*,…, x_n_}*. The *true* agent set *S* is a subset of *U* with *s* = *|S|* elements. We call any *X ⊆ U* a treatment, and the outcome of a treatment is a function

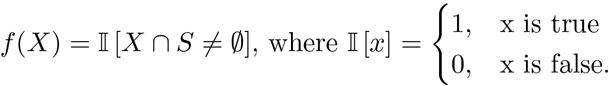

For simplicity, we will refer to treatments that yield an observation of 1 as **positive** outcomes, and those that yield 0 as **negative** outcomes.

We define a **round** as the time spent from setting up a treatment test to collecting the treatment test result. We assume all treatments require an equal amount of time to be tested. We define **efficiency** *w* as the maximal number of parallel treatments that can be tested in a single round. For example, in Figure 1, we can test 4 treatments in a single round, so efficiency *w* = 4.

We define the *sample bound O*(*n, s*) of an algorithm as the upper bound of the total tests required for exactly identifying *S* for *n* candidates. We also define a *Time bound T*(*n, s, w*) as the upper bound of the total rounds needed for exactly identifying *S* for *n* candidates at efficiency *w*.

A **naive algorithm** is to test each candidate in its own treatment. We can exactly identify all agents using *n* tests in *⌈n/w⌉* rounds. Hence, this method has sample bound *O*(*n, s*) = *n* and time bound *T*(*n, s, w*) = *⌈n/w⌉*.

In the following sections, we will describe a novel **group-screening algorithm** that has sample bound 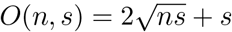 and time bound 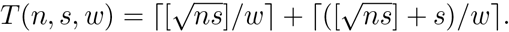

### 2.2 Group-screening

The intuition behind the group-screening method is that by splitting the candidate set to an appropriate number of groups, we may be able to identify a significant portion of non-agents. We call these groups “group-screening treatments”.

If a group-screening treatment has a negative outcome, we can mark all the candidates in this treatment as non-agents. In the next step, we only need to test the candidates from group-screening treatments that have a positive outcome. The size of a group-screening treatment refers the number of candidates pooled together. Group number *b* is the number of group-screening treatments to be tested. In an extreme setting, putting each candidate in its own treatment, where *b* = *n*, recovers the naive method. We present this method in Algorithm 1.

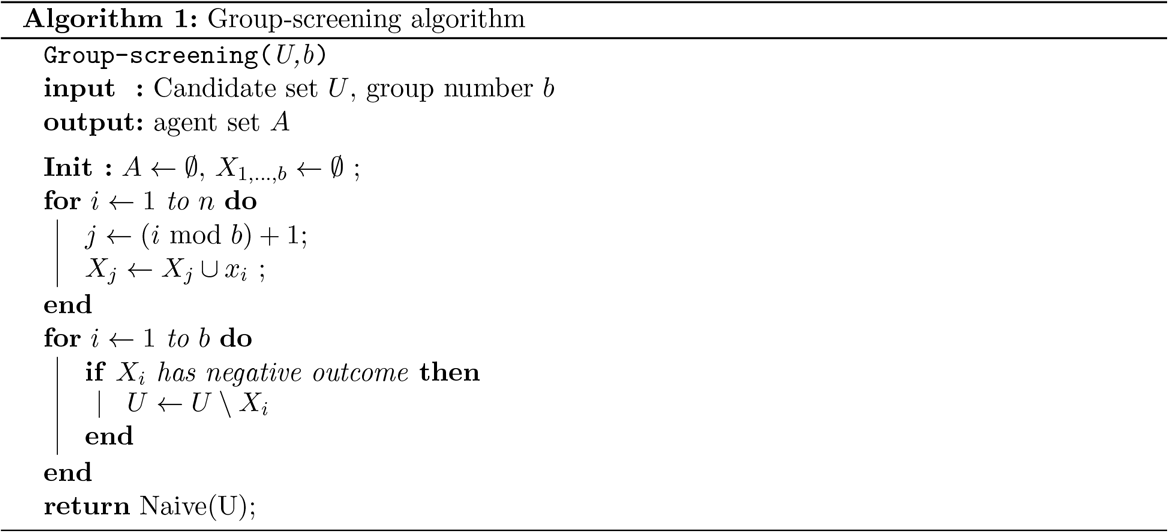

We show two lemmas for group-screening method here:

**Lemma 1.** *Using group-screening for n candidates with b group-screening treatments, if there are at most s agents, Algorithm 1 will require at most* 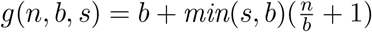 *tests in total.*

*Proof.* If *b ≤ s*, in the worst case, each of the *b* group-screening treatments will have a positive outcome, hence the algorithm will require *n* more tests to find all the agents resulting in *b* + *n* tests in total. If *b > s*, there will be at most *s* group-screening treatments with positive outcomes, and at least *b − s* treatments will have negative outcomes. The algorithm will require 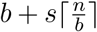 tests in total. Combining both cases, we will require no more than 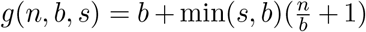 tests.

If *b > s*, we can find *b^*^* that will minimize *g*(*n, b, s*) wrt *b* by taking the derivative of *g* and equating it to zero. This gives 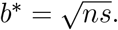 As *b* is an integer, we can take 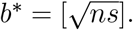 We call *b^*^* the optimized group number, as it minimizes the sample bound *g*(*n, b, s*). By choosing this optimized group number, we can derive the following theorem:

**Theorem 1.** *Using group-screening method on n candidates with b group-screening treatments, if there are at most s agents, Algorithm 1 with* 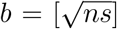 *will require at most* 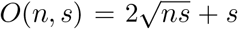 *tests.*

*Proof.* As *n ≥ s*, we have *b^*^ ≥ s*. Therefore, we have: 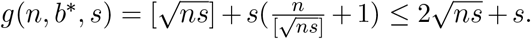 Hence, we can test 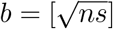 group-screening treatments to identify all agents with no more than 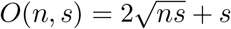 tests.

Under the optimal group number 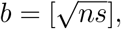 Algorithm 1 requires 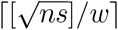 rounds for group-screening step, and at most 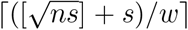 for naive method step. Hence, the upper bound on time is 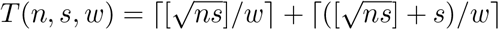 rounds.

### 2.3 Decision rules for methods

The sample bound for the group-screening method can go beyond *n* when *s* is large, which is less efficient than using the naive method. Given *n* candidates with at most *s* agents, we wish to choose the most efficient method. Here, we discuss the decision rules for method selection. The naive method is a deterministic algorithm and will always need to test *n* treatments to identify agents among *n* candidates. Thus, if the sample bound is lower than the naive method’s cost *n*, we can be sure to save tests in the worst case.

We give the following lemmas without proof as they require only simple algebra.

**Lemma 2.** *Given n candidates, if there are at most s agents and 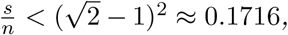 group-screening algorithm (Algorithm 1) will always require less than n tests to identify all of the agents.*

A similar analogy can be conducted for time bound:

**Lemma 3.** *Given n candidates and efficiency w, if there are at most s agents and 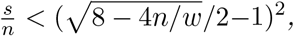 the group-screening algorithm (Algorithm 1) will always need at most ⌈n/w⌉ rounds to identify all of the agents.*

We summarize these features in Table 2.

**Table 2.**
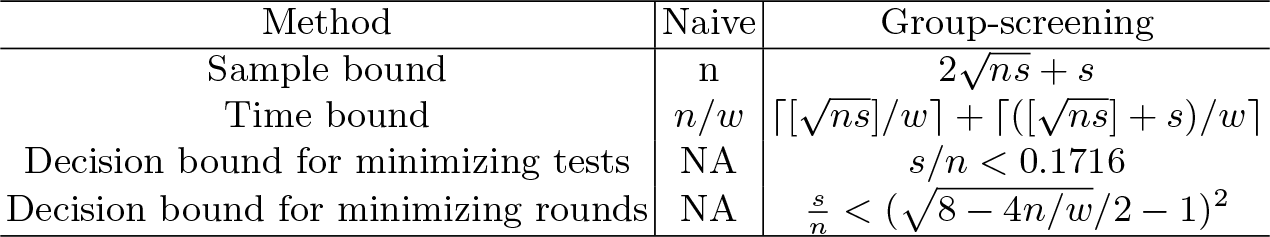
Comparison between naive and group-screening methods

To better illustrate the savings, we introduce the concept of saving ratio. The saving ratio indicates the percentage of tests or rounds that we can save. Let *n^*^* be the actual total tests, we define test saving ratio *Ω_t_* = 1 *− n^*^/n*. Similarly, let *r^*^* be the actual total rounds, we define round saving ratio as *Ω_r_* = 1*−r^*^/⌈n/w⌉*. Using Lemma 2 and 3, we can derive the lower bound for saving ratios using the group-screening method:

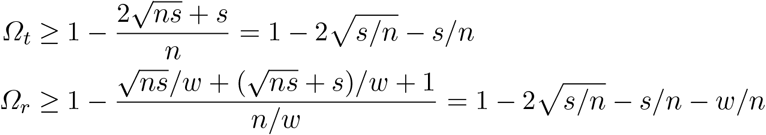

Intuitively, the smaller the true agent number, the more we can save. The relationship between true agent ratio *s/n* and test saving ratio bound is shown in Figure 2 (a).

The round saving ratio bound is a function of both true agent ratio *s/n* and efficiency ratio *w/n*. The smaller the true agent ratio and efficiency ratio are, the more we can save. We showed this relationship in Figure 2 (b).

These bounds provide a reference for how much we can save by using the group-screening method. However, this is a worst case analysis. On average we can save more than this. We showed the worst case bound and empirical savings in Figure 3. The average saving ratio is typically larger than the worst case bound.

**Fig. 2.**
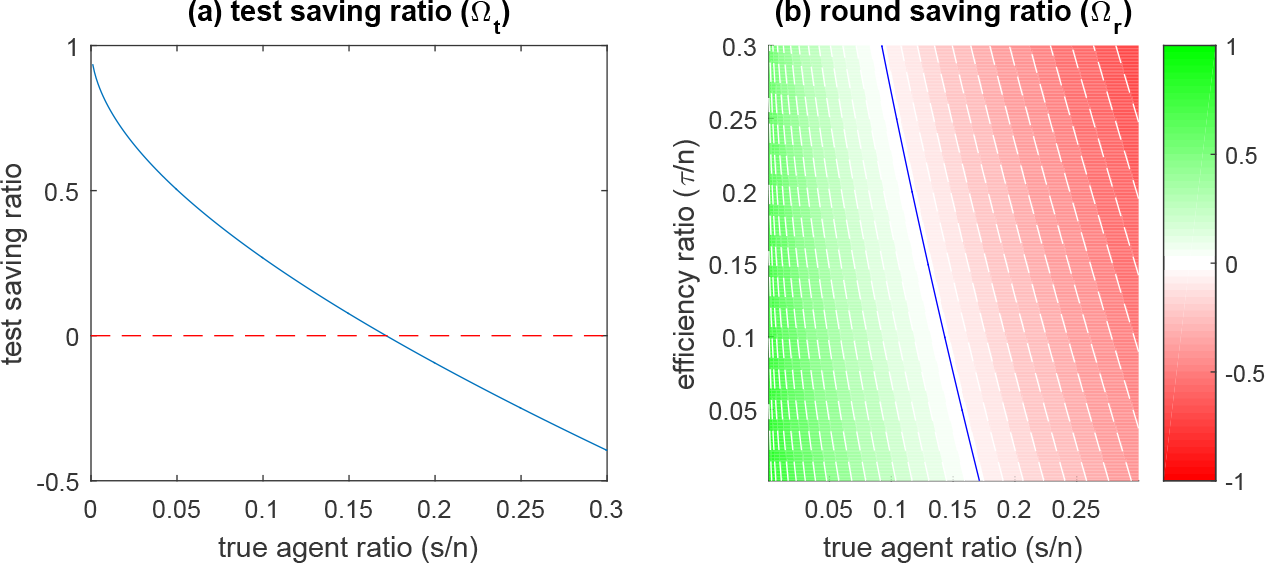
(a) The test saving ratio lower bound as a function of true agent ratio. The blue line indicates the worst case saving ratio under the given true agent ratio. The smaller the true agent ratio, the more tests we can save. The red dashed line indicates the point of no savings (naive method). (b) Round saving ratio bound as a function of efficiency ratio and true agent ratio. The blue line indicates the decision boundary of positive savings in terms of rounds. Green indicates the savings, and red indicates the losses. We expect to save when true agent ratio is low and efficiency ratio is low.

**Fig. 3.**
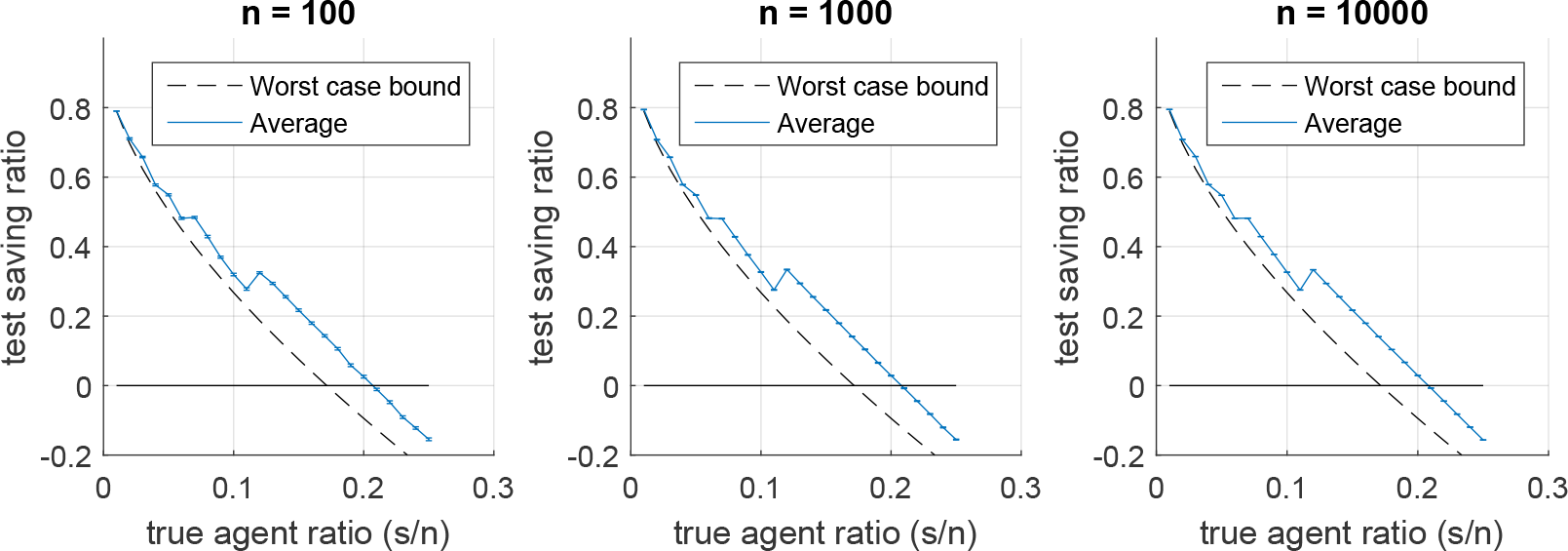
Average test saving ratios for different number of candidates. From left to right, the cases for *n* = 100*, n* = 1000, and *n* = 10000. *Blue* line represents the average test saving ratio over 100 simulations, the error bar indicates the standard error. From simulation we found that the average test saving ratio is generally larger than the worst case bound.

### 2.4 An online-group-screening method

In most cases, we do not know the true number of agents *s* before we start the screening procedure. However, as we test more treatments, we gather more information about the frequency of the true agents. Suppose we have tested *m* treatments, if *k* true agents are identified, we can obtain a reasonable estimate for overall frequency of agents among all candidates using hypergeometric distribution. Here, we provide a probabilistic bound on the total number of agents.

**Theorem 2.** *Given the outcome of m treatments from n candidates and k agent detections, let s be the true agent number in n candidates*, 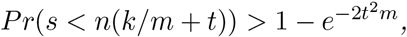 *where t* ≥ 0.

*Proof.* Let 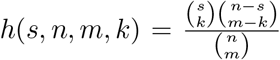 be the hypergeometric distribution probability of detecting *k* agents from randomly chosen *m* candidates among *n* total candidates with *s* agents. Chvatal [10] gives a bound for cumulative hypergeometric distribution tail 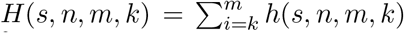 using Hoeffding [11] inequality 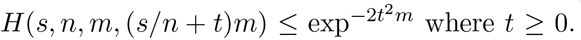 where *t* ≥ 0. This relationship can be rewritten into: 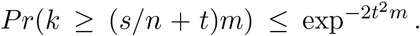 Replacing *s* with the true non-agent number *n − s* and *k* with non-agent detections *q* = *m − k* yields the inequality in Theorem 2 using a sequence of algebraic manipulations on the probability of complementary events.

We can compute the upper bound on the number of agents which will hold with certain probability *a*. This upper bound is given by 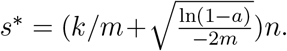 Therefore, the rest of the candidates will contain at most *s^*^ − k* agents with probability more than *a*. The choice of *a* depends on our strategy for making use of existing treatment results. Large *a* is conservative, as it decreases the chance of both loss and saving. Small *a* is risky, as it increases the chance of both loss and saving. In our practical experience, we found that *a* = 0.5 will give us reasonable average savings, and we use *a* = 0.5 in the following analysis.

Given the tolerance for risk, we can set the probability *a* and obtain the corresponding upper bound on true agent number *s*. If the worst case saving ratio under such *s* is positive, we can choose to use the group-screening method for the rest of the candidates in the next round. Otherwise, we will continue to use the naive method for the next round. This online based algorithm is shown in Algorithm 2.

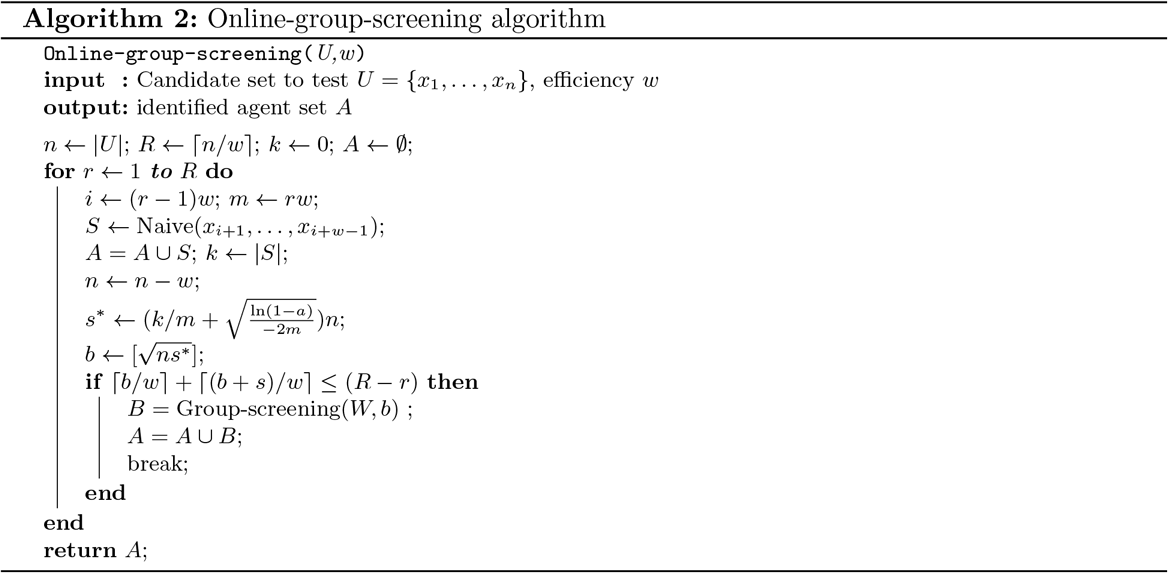

We showed an empirical analysis for the online-group-screening method in Figure 4. From the result, we found that even without knowing the true agent number *s*, our method will still be able to make savings. On the other hand, this method rarely incurs a loss.

### 2.5 Directions regarding the true experiments

Here, we discuss some practical concerns in true experiments with regard to the online-group-screening method.

**Fig. 4.**
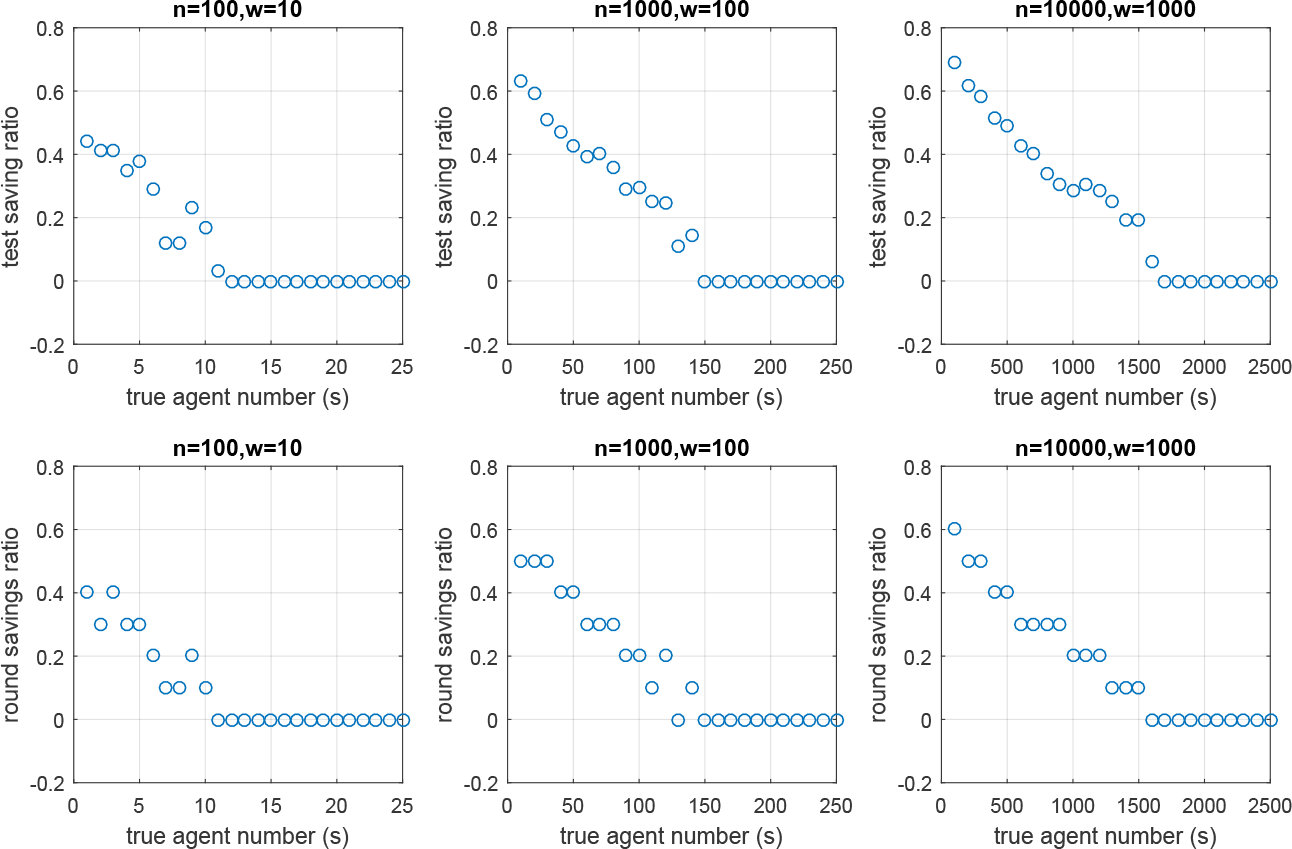
An empirical analysis for the online-group-screening method. We ran 100 simulations for different settings of *n, w*, where true agent ratio is uniformly generated. The first row is the test saving ratio. The second row is the round saving ratio. We only show the result for the first 25% true agent ratio, as the rest 75% have 0 saving ratios in all the settings. No loss has occurred in all 100 simulations for all the settings.

*Candidate limit per treatment.* Many biological or chemical experimental settings naturally limit the number of candidates in a single treatment. Mixing too many candidates may dilute the agents below the threshold required for their effects. Given *n* candidates and candidate limit per treatment set as *τ*, at least 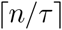 groups are required. We can change the Algorithm 2 by modifying 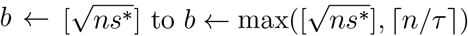 to satisfy this limit.

*Grouping preference* In real experiments, we generally have side information about candidates before we start the screening procedure. Candidates believed to have similar effects should be placed in the same treatment. This design increases the chances that agents are concentrated in a single treatment.

We provide a Matlab implementation of this method: https://bitbucket.org/clingsz/groupscreening/src/.

## 3 Case study for a high-throughput screening experiment

Our method performs well in settings where total candidate number is very large and true agents are very few. This is usually the case for high-throughput screening experiments. We investigated our method on a specific problem from Ma *et al.* [2], where 50,000 chemically diverse compounds were screened for inhibition of cAMP/flavone-stimulated Cl*^−^* transport in epithelial cells expressing cystic fibrosis transmembrane conductance regulator (CFTR), which has a strong relationship with secretory diarrhea.

In this experiment, candidates are *n* = 50, 000 chemical compounds. *s* = 7 compounds with significant effect – the true agents – were found using the high-throughput screening method. Each round, 40-60 96-well plates can be analyzed in parallel. Hence, the work efficiency is *w* = 5760. To fully analyze all 50,000 candidates, the naive method takes 9 rounds and 50,000 tests in total. By knowing *s* = 7, the group-screening algorithm showed that by randomly dividing all the candidates into 592 group-screening treatments, where each treatment contains at most 85 candidates, in the worst case we can identify all the true agents using 2 rounds and 1187 tests in total.

If we have no prior information about *s*, we can use the online-group-screening method by utilizing the results acquired from the first round. Our algorithm decided to randomly split all the rest of the candidates into 4024 group-screening treatments, with at most 11 candidates in each treatment. Finally, we can find all true agents using 3 rounds and 9828 tests, a five-fold saving in test effort. We summarized these savings and compared with the naive method in Table 3.

**Table 3.**
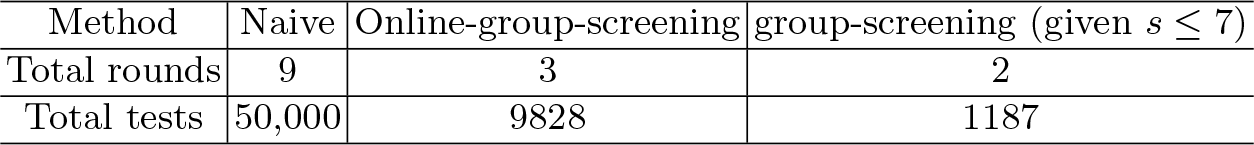
Comparison between naive and online-group-screening methods on HTS

If the maximal number of candidates is limited in each treatment, the online-group-screening method can still save tests. We showed the total number of tests and rounds required under different limitations in Figure 5. Even if at most 2 candidates are allowed in each treatment, we can save both time and tests by almost 30%.

**Fig. 5.**
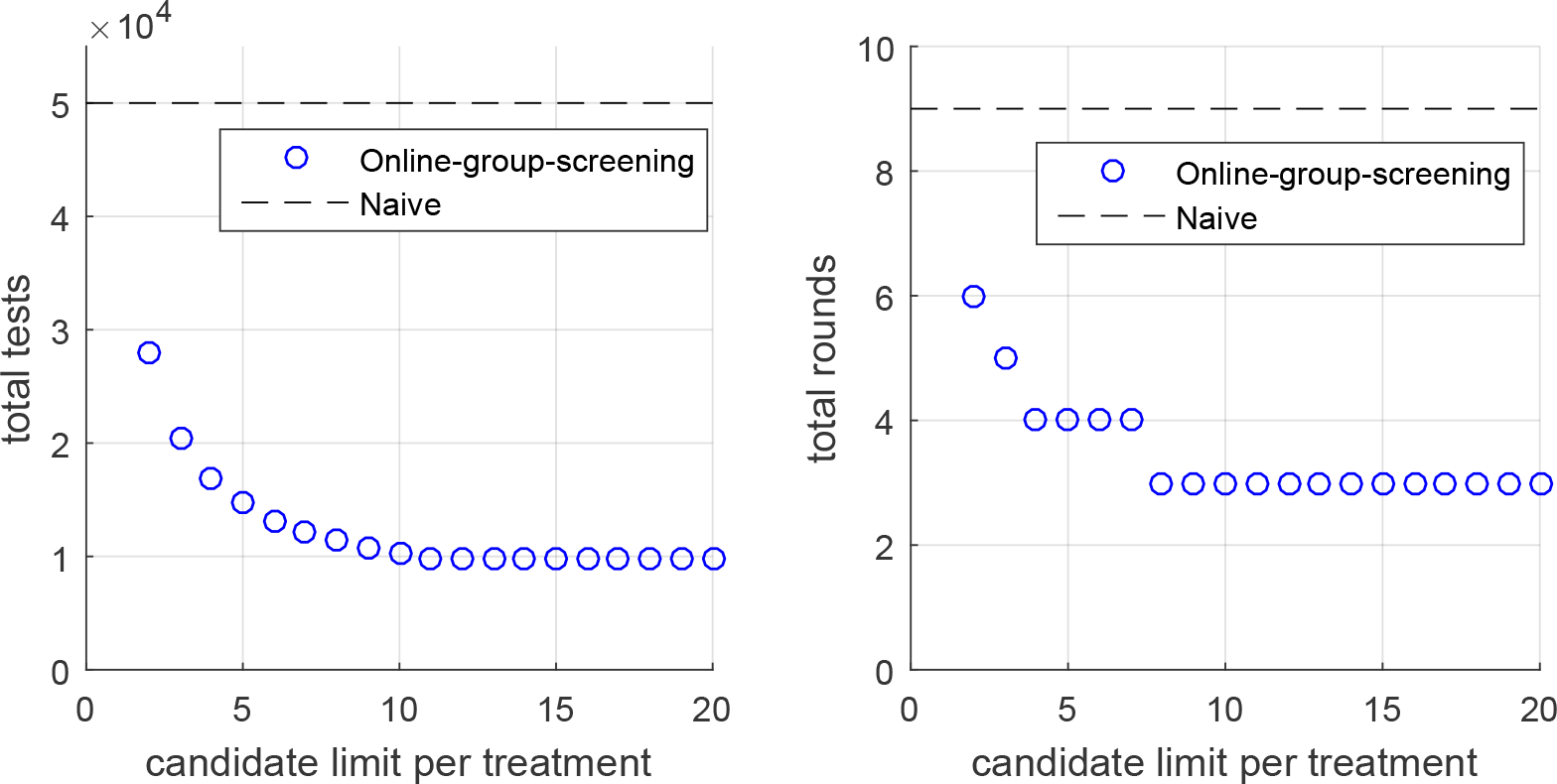
Total tests and rounds needed for the online-group-screening method when candidate number in each treatment is limited. Left: total tests needed. Right: total rounds needed. Blue circles represent the result of online-group-screening method, black dashed line represents the total tests and rounds required by naive method.

## 4 Extensions

We can slightly modify our problem specifications to solve other variants. Next, we show 2 variant problems to illustrate the adaptability of our framework: essential set identification and agent identification for multiple phenotypes.

### 4.1 Essential set identification

Identifying the essential set is a common problem in biology. For example, in order to understand the molecular and biological functions of genes in a cell, Hutchison *et al.* [12] built a minimal genome by including only the essential genes in bacteria. Discovering exact genes that are essential for bacterial survival and growth is a prerequisite for successful constructing such minimal genome. A straightforward naive method is to exclude each gene one by one from the whole genome to test whether it is essential. Suppose we have *n* candidate genes *U* = *{x*_1_*, x*_2_*,…, x_n_}*. The essential set *S* is a subset of *U* with *s* = *|S|* elements. We refer to any *X ⊆ U* as a treatment, where we construct a bacteria without any of the candidates in *X* and let the bacteria grow. The outcome of a treatment is a function 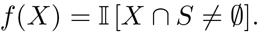

In this function, if the bacteria does not grow when **excluding** candidates in *X*, then *f* (*X*) = 1. This indicates that some of the genes in *X* are essential. Conversely, if the bacteria grow, *f* (*X*) = 0, then none of the candidate compounds in *X* are essential since this exclusion does not affect the bacteria growth. The essential set identification problem can be mapped onto our framework of agent identification.

### 4.2 Agent identification for multiple phenotypes

Causal factors may drive multiple phenotypes. For example, plant growth promoting bacteria can influence plant height, flowering time, number of flowers, or the root architecture [13–16]. When screening for plant growth promoting traits of bacteria, the expected phenotype is often unknown a-priori. Here, we show how our framework can be adapted to the problem of agent identification for multiple phenotypes.

Given measurements of *v* phenotypes and unknown true agent sets *S*_1_*, S*_2_*,…, S_v_* responsible for the corresponding phenotypes, we define the outcome of a treatment for the *i*th phenotype as: 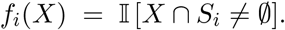 Therefore, f(*X*) = [*f*_1_*, f*_2_*,…, f_v_*]*^T^* is a vector that represents the outcome for *v* phenotypes. For this problem, a naive method can still identify all the *v* agent sets using *n* treatments by testing each candidate one by one. Here, we define the global agent set *S* = *S*_1_ ⋃ *S*_2_ ⋃ *· · · ⋃ S_v_*, and treatment outcome function 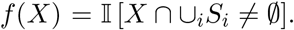 Effectively, a treatment has positive outcome if any phenotype is affected by an agent, and negative if no phenotype was affected.

Thus, this problem is equivalent to the agent identification for a single abstract phenotype defined as a logical OR of the multiple phenotypes under study.

## 5 Discussion

Efficient identification of agents among a large number of candidates is a challenging problem in biology and medicine. In this paper, we introduced an efficient online-group-screening framework that can 1) automatically decide the optimal strategy based on the outcome of a small set of pilot treatments, 2) requires less experimental rounds and total tests than the exhaustive method even in the worst case. We showed both the worst case savings and empirical average savings under different settings. Our method adapted to an existing dataset gives the expected time and cost savings. We also showed that this framework can be easily modified to solve related problems. We believe this method will make agent identification problems less expensive and much more efficient, and thus increase the pace of discovery.

*Future work* This approach can be further applied to the identification of more complex causal mechanisms, for example, involving interacting agents. If true interactions between candidates are few, we can still expect to save cost and time.

